# A novel function for Rab proteins during maturation of secretory granules

**DOI:** 10.1101/2021.03.01.433309

**Authors:** Sarah D. Neuman, Annika R. Lee, Jane E. Selegue, Amy T. Cavanagh, Arash Bashirullah

## Abstract

Regulated exocytosis is an essential process whereby professional secretory cells synthesize and secrete specific cargo proteins in a stimulus-dependent manner. Cargo-containing secretory granules are synthesized in the *trans-Golgi* Network (TGN); after budding from the TGN, granules undergo many modifications, including a dramatic increase in size. These changes occur during a poorly understood process called secretory granule maturation. Here we leverage the professional secretory cells of the *Drosophila* larval salivary glands as a model system to characterize a novel and unexpected role for Rab GTPases during secretory granule maturation. We find that secretory granules in the larval salivary glands increase in size ~300-fold between biogenesis and release, and loss of *Rab1* or *Rab11* dramatically reduces granule size. Surprisingly, we find that Rab1 and Rab11 protein localize to secretory granule membranes. Rab11 associates with granule membranes throughout the maturation process, and Rab11 is required for recruitment of Rab1. In turn, Rab1 associates specifically with immature secretory granules and drives granule growth. In addition to their roles in granule growth, both *Rab1* and *Rab11* appear to have additional roles during exocytosis; *Rab11* function is necessary for exocytosis, while the presence of *Rab1* on immature granules may prevent precocious exocytosis. Overall, these results highlight a new and unexpected role for Rab GTPases in secretory granule maturation.

## INTRODUCTION

Regulated exocytosis begins at the *trans-Golgi* network (TGN); here, cargo proteins destined for exocytosis are packaged into specialized organelles called secretory granules. After budding from the TGN, nascent granules undergo many modifications in a process called secretory granule maturation (Kögel and Gerdes, 2010a; Bonnemaison et al., 2013). One of the conspicuous modifications that occurs during maturation is an increase in secretory granule size as immature granules fuse with one another (Kögel and Gerdes, 2010a; Bonnemaison et al., 2013; Hammel et al., 2010). Secretory granule ‘growth’ has been described in a variety of mammalian professional secretory cells, including pancreatic acinar cells (Lew et al., 1994; Weintraub et al., 1992), pancreatic β-cells (Du et al., 2016), mast cells (Hammel et al., 1983, 1985), pituitary mammotrophs (Smith et al., 1966; Farquhar et al., 1978), and PC12 cells (derived from adrenal chromaffin cells) (Ahras et al., 2006; Urbé et al., 1998; Wendler et al., 2001). Although this phenomenon has frequently been observed, the molecular mechanisms that regulate secretory granule growth remain poorly understood.

Rab proteins are small GTPases that play a role in regulating nearly every aspect of intracellular trafficking, including regulated exocytosis. These proteins are highly conserved throughout eukaryotes, although the number of Rabs increases with the complexity of the organism; the yeast genome contains 11 Rab proteins, while the human genome contains at least 60 (Pfeffer, 2017; Li and Marlin, 2015; Homma et al., 2021). Rab proteins are prenylated at their C-terminus, allowing for membrane localization (Kinsella and Maltese, 1991; Leung et al., 2006; Rossi et al., 1991; Khosravi-Far et al., 1991). Inactive Rab proteins are bound to GDP and GDP-dissociation inhibitor (GDI) and localized to the cytoplasm; upon approaching their target membrane, GDI is displaced by GDI displacement factor (GDF), allowing membrane insertion of the Rab protein (Li and Marlin, 2015; Sasaki et al., 1990; Pfeffer et al., 1995; Müller and Goody, 2018; Sivars et al., 2003). The Rab protein then interacts with a cognate guanine nucleotide exchange factor (GEF), which facilitates the replacement of GDP for GTP, activating the Rab (Blümer et al., 2013; Müller and Goody, 2018; Ullrich et al., 1994; Soldati et al., 1994; Barr and Lambright, 2010). GTP-bound Rabs physically interact with their effector proteins to regulate membrane trafficking, including vesicle budding, movement, and fusion, among other functions (Homma et al., 2021; Li and Marlin, 2015). Rab proteins are inactivated via interaction with GTPase activating proteins (GAPs) which aid in catalyzing the hydrolysis of GTP to GDP, allowing the Rab to be removed from the membrane via interaction with GDI (Ullrich et al., 1993; Soldati et al., 1993; Garrett et al., 1993; Müller and Goody, 2018; Barr and Lambright, 2010). Each Rab protein is localized to specific membrane compartments, allowing these small GTPases to direct and orchestrate many of the complex trafficking events required for regulated exocytosis. Rab3D and Rab27A have previously been reported to localize to secretory granule membranes in mammalian neuroendocrine and endothelial cells, respectively, where they may play a functional role in regulating secretory granule membrane and cargo remodeling during maturation (Kögel and Gerdes, 2010b; Kögel et al., 2013; Hannah et al., 2003). However, a role for other Rab proteins in secretory granule maturation in different cell types has not been described.

The *Drosophila* larval salivary glands are composed of professional secretory cells that synthesize and secrete massive quantities of mucin-like “glue” proteins via regulated exocytosis (Korge, 1977; Beckendorf and Kafatos, 1976). Mucin biogenesis and secretion are developmentally controlled. Biogenesis begins at the mid-third instar larval transition, and exocytosis begins about 4 h before the onset of metamorphosis, 24 h later (Biyasheva et al., 2001; Kang et al., 2017; Beckendorf and Kafatos, 1976). Once exocytosis is complete, the mucins are expelled out of the lumen of the salivary glands and onto the surface of the animal, where they act like glue, allowing the puparium to adhere to a solid surface during metamorphosis (Biyasheva et al., 2001; Kang et al., 2017). Like secretory granules in many types of mammalian cells, these mucin-containing granules undergo a maturation process, increasing in size over time (Niemeyer and Schwarz, 2000; Reynolds et al., 2019; Neuman and Bashirullah, 2018). Because mucin production is a developmentally controlled process consisting of a single round of biogenesis, maturation, and exocytosis, we are able to view the events that occur during maturation with a high degree of temporal resolution. Moreover, fluorescent tagging of mucins (*Sgs3-GFP* or *Sgs3-DsRED*) (Biyasheva et al., 2001; Costantino et al., 2008) allows us to view these events in living tissue in real-time.

Here, we have leveraged mucin secretion in the larval salivary glands as a model system to examine the role of Rab proteins in secretory granule maturation. We screened *Drosophila* Rab proteins using stage- and tissue-specific RNAi knockdown for defects in secretory granule size. This screen demonstrated that RNAi knockdown of *Rab1* and *Rab11* resulted in a striking small granule phenotype. Surprisingly, we found that Rab1 and Rab11 protein localized to secretory granule membranes; Rab11 was present on granule membranes from biogenesis through secretion, while Rab1 was only present on granule membranes during the process of maturation. We also demonstrate that Rab1 and Rab11 localization to granule membranes requires GTP binding, and that Rab11 is required for recruitment of Rab1. *Rab11* function is required for both maturation and exocytosis, as loss of *Rab11* results in secretion defects and impairs the ability of granules to dock and fuse with the apical membrane. Finally, surprisingly, we found that small granules lacking Rab1 are able to be secreted, suggesting that *Rab1* may have a role in preventing precocious secretion of immature granules. Overall, these results highlight a previously unknown role for Rab proteins on secretory granule membranes during the process of secretory granule maturation and provide new insights into the role of maturation during regulated exocytosis.

## RESULTS

### Secretory granules dramatically increase in size from biogenesis to secretion

The biogenesis of mucin-containing secretory granules in the *Drosophila* larval salivary glands begins when animals reach the mid-third instar larval transition and ends with secretion at the onset of metamorphosis (puparium formation) 24 h later (Beckendorf and Kafatos, 1976; Biyasheva et al., 2001). Previous work has shown that mucin-containing granules undergo a maturation process, increasing in size by fusing with one another (Neuman and Bashirullah, 2018; Niemeyer and Schwarz, 2000; Reynolds et al., 2019). To further characterize and quantify changes in secretory granule size that occur during this 24 h period from biogenesis to secretion, we synchronized animals at the onset of mucin biogenesis and used confocal microscopy to analyze mucin granule size in 4 h windows spanning this time period. We defined granules in the 0-4 h timepoint as ‘nascent granules,’ granules in the 4-20 h period as ‘immature granules,’ and granules in the 20-24 h period (when exocytosis has begun) as ‘mature granules’ (Fig. 1A). We observed that granules dramatically increased in size, with nearly a 300-fold increase in overall volume when comparing nascent vs. mature granules (Fig. 1A, C). Interestingly, the single largest increase in volume occurred between the 0-4 h and 4-8 h timepoints, with ~18-fold change in average volume, indicating that the granules doubled in size every 1 h (Fig. 1C). Thereafter, the secretory granules doubled in size about every 4 h, with a total of ~15-fold change in average volume between the 4-8 h and 20-24 h timepoints (Fig. 1C). This dramatic difference in growth rate between nascent granules and immature granules suggests that there may be independent regulatory mechanisms mediating each of these stages of granule growth.

**Figure 1.**
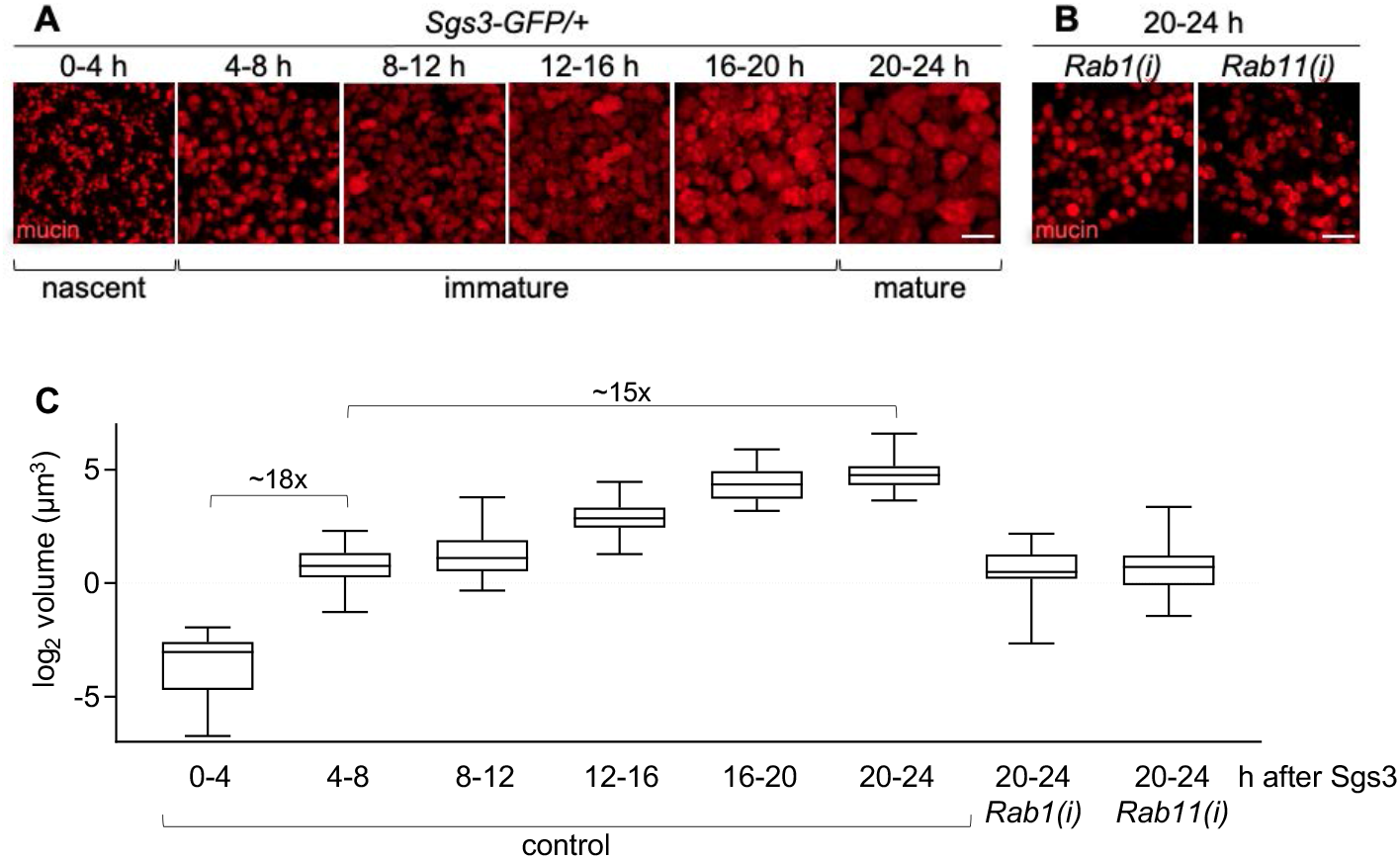
Analysis of secretory granule volume during maturation. **(A)** Live-cell imaging of mucins (red) from staged salivary glands spanning from the onset of biogenesis (0-4 h) to the onset of exocytosis (20-24 h). Timepoints are shown in h after Sgs3. Images shown are a maximum intensity projection of ten optical slices from z-stacks comprising 24-32 total slices at a step size of 0.28 μm. **(B)** Live-cell imaging of mucin granules (red) at the 20-24 h after Sgs3 timepoint in salivary glands expressing *Rab1-* or *Rab11-RNAi*. Full genotypes: *Sgs3-GFP/UAS-Rab1-RNAi; Sgs3>/+* and *UAS-Rab11-RNAi/+; Sgs3-GFP/+; Sgs3>/+*. Scale bars: 5 μm. **(C)** Box and whisker plot showing the volume of secretory granules from staged salivary glands. Z-stack imaging data was used to identify the medial cross-section of each granule for quantification. *x*-axis reflects the developmental stages and genotypes analyzed; *y*-axis shows log_2_-transformed volumes in μm^3^. Boxes outlines the 25th to 75th percentiles; middle line indicates the median. Whiskers extend to the minimum and maximum values. *n*=50 granules per timepoint/genotype.

### An RNAi screen for Rab proteins that regulate secretory granule size

The *Drosophila* genome contains 33 annotated Rab proteins (Zhang et al., 2007). To test if any of these Rabs play a role in regulating secretory granule size, we conducted a salivary gland-specific RNAi screen and looked for a reduction in mature granule size. We used the stage- and tissue-specific *Sgs3-GAL4* driver for this screen (Cherbas et al., 2003); since GAL4 expression is driven by the same promoter as mucins, we were able to examine specific effects on mucin granules without affecting development or earlier functions of the larval salivary glands. We tested each of the 30 annotated *Drosophila* Rab proteins that had one or more RNAi lines publicly available. This screen identified *Rab1* and *Rab11* as high-confidence hits with a dramatic reduction in secretory granule size validated by multiple independent RNAi lines (Table S1, Fig. 1B, C). We also identified *Rab3, Rab5, Rab10*, and *Rab27* as lower-confidence hits with a moderate reduction in granule size that was only observed in a single RNAi line (Table S1). qPCR analysis in dissected salivary glands confirmed that *Rab1-RNAi* and *Rab11-RNAi* significantly reduced *Rab1* and *Rab11* expression levels, respectively (Fig. S1). The terminal size of mucin granules in salivary glands expressing *Rab1-* or *Rab11-RNAi* was similar to that of controls at the 4-8 h timepoint (Fig. 1C), indicating that disruption of *Rab1* or *Rab11* function severely inhibits mucin granule growth. These screen results highlight a novel role for Rab proteins in the regulation of secretory granule size.

### Rab1 and Rab11 localize to secretory granule membranes

Many Rab proteins localize to specific organelle membranes, and this specificity in localization allows them to carry out their roles in intracellular trafficking (Li and Marlin, 2015; Homma et al., 2021; Zhen and Stenmark, 2015). To begin to gain an understanding of how Rab proteins regulate mucin granule size, we used a collection of endogenously regulated, EYFP-tagged Rab proteins (Dunst et al., 2015) to assess the subcellular localization patterns of our screen hits in salivary glands during the 24 h time period spanning mucin biogenesis through secretion. Surprisingly, we observed that both Rab1 and Rab11 were enriched on secretory granule membranes. Rab11 appeared to be enriched on secretory granule membranes throughout the entire 24 h period from biogenesis through secretion (Fig. 2A, C). Rab11 has typically been reported to localize to the Golgi and to recycling endosomes (Zhen and Stenmark, 2015; Ullrich et al., 1996; Calhoun and Goldenring, 1996; Welz et al., 2014; Urbé et al., 1993), and we did observe co-localization of Rab11 with a pan-Golgi marker, as well as small Rab11-positive puncta that are likely recycling endosomes (Fig. S2B), indicating that Rab11 localizes both to the expected organelles and to secretory granule membranes. Rab1, in contrast, exhibited stagespecific localization to granule membranes; Rab1 was absent from both nascent (0-4 h) and mature (20-24 h) granules but present on nearly all granules at the 4-8 h and 8-12 h timepoints and present on some granules at the 12-16 h and 16-20 h timepoints (Fig. 2A, B). Rab1 is typically thought to localize to endoplasmic reticulum exit sites (ERES), where it plays a role in promoting anterograde ER-to-Golgi trafficking (Zhen and Stenmark, 2015; Plutner et al., 1991; Segev et al., 1988; Schmitt et al., 1988). We observed co-localization of Rab1 with Sec31 (Fig. S2A), a component of the COPII coat protein complex and marker for ERES (Förster et al., 2010), indicating that Rab1 also localizes to ERES in salivary glands. In contrast, our lower-confidence screen hits did not appear to localize to secretory granule membranes (Fig. S3), suggesting that Rab1 and Rab11 may play a granule-autonomous role in regulating secretory granule size.

**Figure 2.**
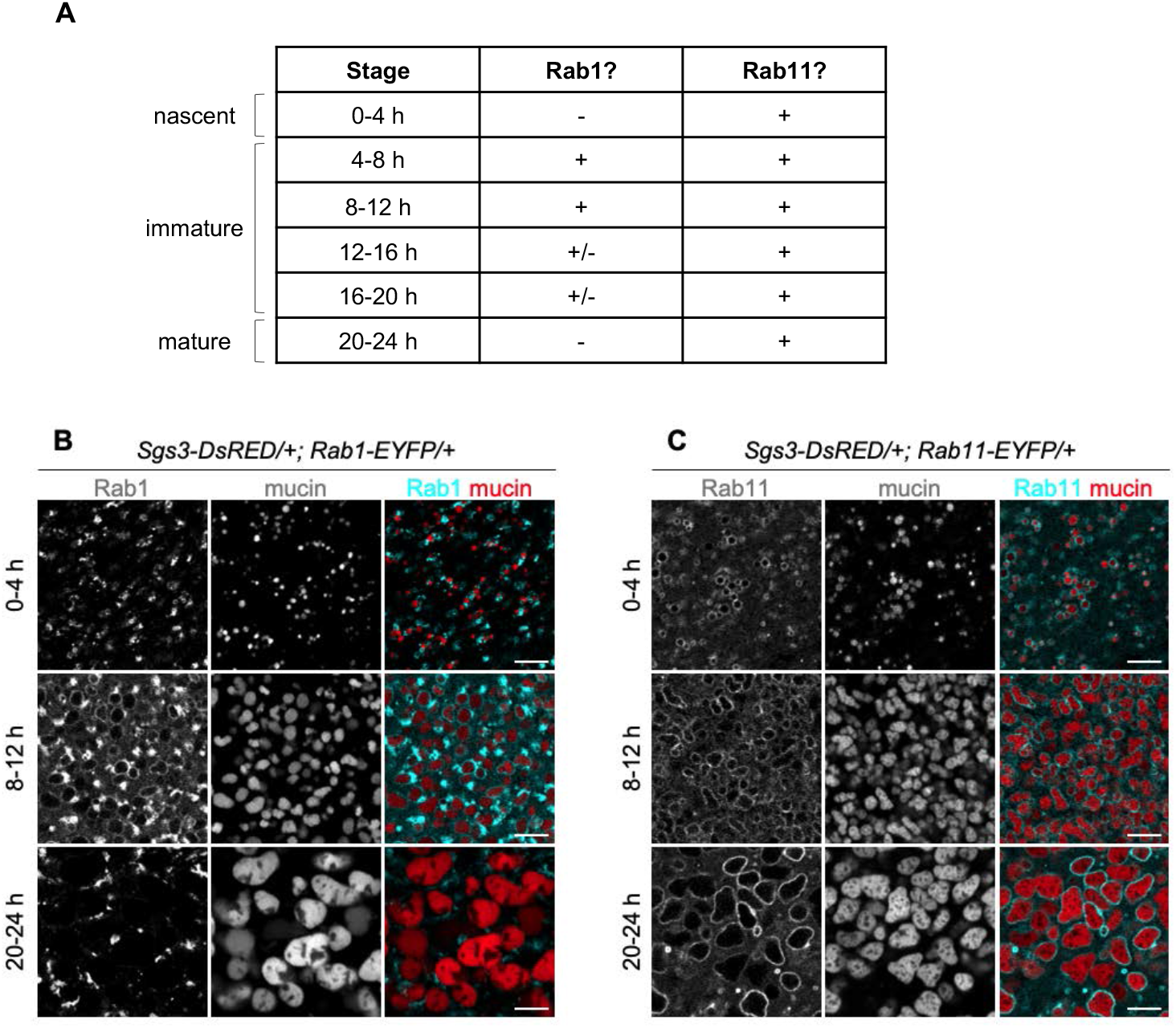
Association of Rab proteins with secretory granules during maturation. **(A)** Summary table showing the presence of Rab1 and Rab11 on secretory granule membranes in staged salivary glands. + indicates the protein is present on most granule membranes, +/- indicates the protein is present on some granule membranes, and – indicates that the protein is not present on granule membranes. Developmental stages are shown in h after Sgs3. *n≥10* glands imaged per timepoint. **(B)** Live-cell imaging analysis of Rab1-EYFP (cyan) localization with mucins (red). Rab1 is absent from granule membranes at 0-4 h after Sgs3, present at 8-12 h after Sgs3, and absent at 20-24 h after Sgs3. Images shown are a single slice from a z-stack comprising three slices at a 0.30 μm step size. **(C)** Live-cell imaging analysis of Rab11-EYFP (cyan) localization with mucins (red). Rab11 is present on granule membranes at 0-4, 8-12, and 20-24 h after Sgs3. Images shown are a single slice from a z-stack comprising ten slices at a 0.31 μm step size. Scale bars: 5 μm.

### GTP binding is required for Rab1 and Rab11 localization and function in maturation

Rab protein function is dependent on binding to GTP; only GTP-bound Rab proteins can recruit their effector proteins to carry out their biological functions (Li and Marlin, 2015; Homma et al., 2021). To examine the role of GTP binding in *Rab1* and *Rab11* function during mucin granule maturation, we overexpressed YFP-tagged dominant negative *Rab1* and *Rab11* specifically in the salivary glands using the *Sgs3-GAL4* driver. These dominant negative constructs contain a point mutation that changes a serine in the GTP binding domain to an asparagine, effectively preventing association with GTP (Zhang et al., 2007). Localization analysis showed that Rab1^DN^-YFP was absent from secretory granule membranes at the 8-12 h timepoint (Fig. 3B), when wildtype Rab1-EFYP is highly enriched on secretory granule membranes (Fig. 3A). Similarly, Rab11^DN^-YFP did not appear enriched on secretory granule membranes at the 8-12 h timepoint (compare Fig. 3C vs. D). These results suggest that GTP binding is required for localization of Rab1 and Rab11 to secretory granule membranes. We did find that expression of the Rab11^DN^-YFP construct was delayed compared to Rab1^DN^-EYFP; we observed robust expression of Rab1^DN^-YFP by ~4 h after Sgs3 but did not see significant expression of Rab11^DN^-YFP until ~8 h after Sgs3. When examining the effect of *Rab1^DN^* and *Rab11^DN^* overexpression on secretory granule size, we found that salivary gland-specific overexpression of both of these proteins resulted in a substantial decrease in secretory granule size at the 20-24 h timepoint (Fig. 3E). However, the phenotype appeared slightly weaker and more variable in *Rab11^DN^*-expressing glands, likely due to delayed expression of this transgenic construct. Overall, this data validates our RNAi screen results and indicates that GTP binding is required for *Rab1* and *Rab11* function during mucin granule maturation.

**Figure 3.**
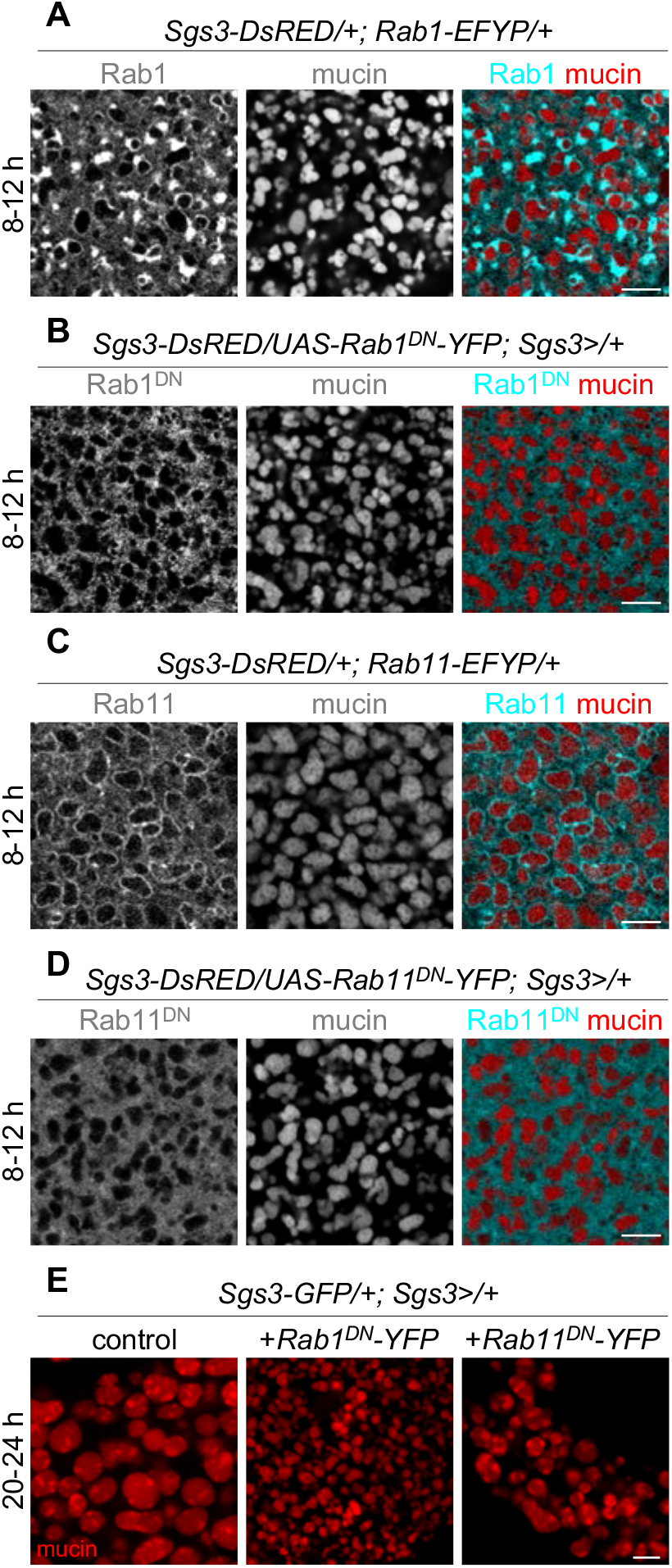
Dominant-negative Rab1 and Rab11 do not localize to secretory granule membranes. **(A)** Live-cell imaging of Rab1-EYFP (cyan) and mucin (red) localization at 8-12 h after Sgs3. Rab1 is present around granule membranes at this timepoint. Image shown is a single slice from a z-stack comprising three slices at a step size of 0.30 μm. **(B)** Live-cell imaging of Rab1 ^DN^-YFP (cyan) and mucin (red) localization at 8-12 h after Sgs3. Rab1^DN^ is not enriched around granule membranes at this timepoint. Image shown is a single slice from a z-stack comprising 12 slices at a 0.31 μm step size. **(C)** Live-cell imaging of Rab11-EYFP (cyan) and mucin (red) localization at 8-12 h after Sgs3. Rab11 is present around granule membranes at this timepoint. Image shown is a single slice from a z-stack comprising ten slices at a step size of 0.31 μm. **(D)** Live-cell imaging of Rab11^DN^-YFP (cyan) and mucin (red) localization at 8-12 h after Sgs3. Rab11^DN^ is not enriched around granule membranes at this timepoint. Image shown is a single slice from a z-stack comprising ten slices at a 0.31 μm step size. **(E)** Livecell imaging of mucin granules (red) in control and Rab1^DN^- and Rab11^DN^-expressing glands at 20-24 h after Sgs3. Mucin granules are reduced in size in Rab1^DN^- and Rab11^DN^-expressing glands. Full genotypes: control- *Sgs3-GFP/+; Sgs3>/+*. Rab1^DN^: *Sgs-GFP/UAS-Rab1^DN^-YFP; Sgs3>/+*. Rab11^DN^: *Sgs3-GFP/UAS-Rab11^DN^-YFP; Sgs3>/+*. Scale bars: 5 μm.

### Rab11 is required for Rab1 localization to secretory granule membranes

Many Rabs function sequentially to regulate membrane trafficking; for example, Rab5 and its effectors are required for Rab7 recruitment to endosomal membranes, regulating the transition from an early endosome to a late endosome (Wandinger-Ness and Zerial, 2014). To test if *Rab1* and *Rab11* function in this manner on secretory granule membranes, we tested if RNAi knockdown of one of these Rabs affected localization of the other. Since Rab11 was present on the membranes of nascent secretory granules (0-4 h timepoint) while Rab1 was not, we first tested if *Rab11* knockdown affected localization of Rab1 protein. Strikingly, we found that Rab1 was largely absent from secretory granule membranes at 4-8 h after Sgs3 upon knockdown of *Rab11* (compare Fig. 4A vs. B), with a significant reduction in the percentage of granules with Rab1 on their membrane at this timepoint (Fig. 4C). Additionally, in cases where secretory granules did have Rab1 present, the amount of Rab1 was dramatically reduced upon loss of *Rab11* (Fig. 4A, B). qPCR analysis of dissected salivary glands showed that *Rab11-RNAi* did not significantly alter *Rab1* mRNA expression levels (Fig. S1), suggesting that loss of Rab11 specifically affects Rab1 localization, not expression. In contrast, RNAi knockdown of *Rab1* did not alter *Rab11* expression levels or localization patterns (Fig. S1, S4A, B). Taken together, these results suggest that Rab11 is required for Rab1 recruitment to the membrane of immature secretory granules.

**Figure 4.**
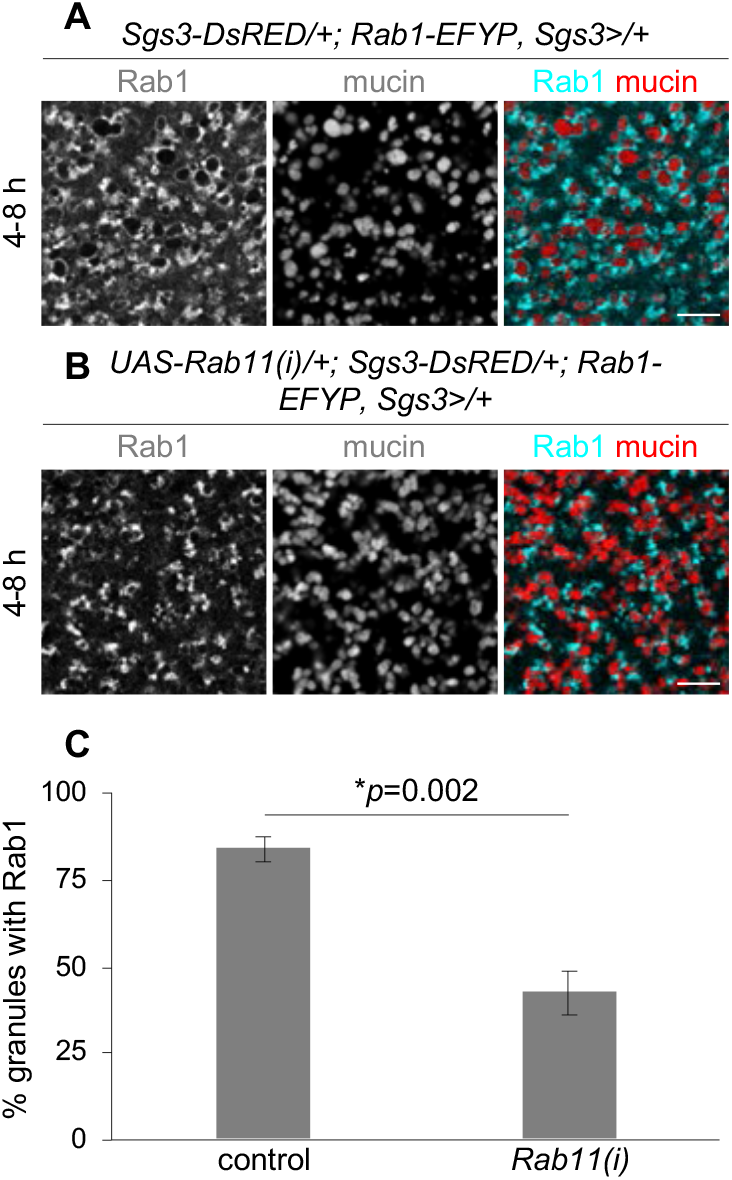
Rab11 is required for Rab1 localization to mucin granule membranes. **(A)** Live-cell imaging of Rab1-EFYP (cyan) and mucin (red) localization at 4-8 h after Sgs3. Rab1 is enriched around granule membranes at this timepoint. Image shown is a single slice from a z-stack comprising six slices at a 0.31 μm step size. **(B)** Live-cell imaging of Rab1-EYFP (cyan) and mucin (red) localization upon salivary gland-specific expression of *Rab11-RNAi* at 4-8 h after Sgs3. Rab1 no longer strongly localizes to mucin granule membranes. Image shown is a single slice from a z-stack comprising six slices at a 0.31 μm step size. Scale bars: 5 μm. **(C)** Quantification of the percentage of granules surrounded by Rab1 shows that there is a significant reduction in *Rab11-RNAi*-expressing glands compared to controls at 4-8 h after Sgs3. Genotypes are as shown for (A) and (B) above. Graph shows mean +/- s.d. from cropped images of three cells from three separate salivary glands. Total *n* of granules counted: control, 393; *Rab11-RNAi*, 433. Statistics calculated by unpaired, two-tailed *t*-test.

### Rab11 function is required for exocytosis

Interestingly, in control cells, we observed that Rab11 protein continued to associate with the membrane of secretory granules during the process of exocytosis, and Rab11 was deposited on the apical plasma membrane after completion of granule fusion (Movie 1, Fig. 5A), suggesting that Rab11 may play a functional role in exocytosis. Some animals homozygous for the hypomorphic *Rab11^93Bi^* mutation (Giansanti et al., 2007; Eisenberg et al., 1990; Jankovics et al., 2001) survive to enter metamorphosis, making it possible to test for exocytosis defects in this mutant background. All mucin proteins had been secreted by the onset of metamorphosis in the salivary glands of control animals; however, *Rab11^93Bi^* mutant salivary glands still contained a significant amount of mucin granules (Fig. 5B), indicating that there were exocytosis defects upon loss of *Rab11* function. Notably, the un-secreted granules were quite small, further confirming a functional role for *Rab11* in regulating secretory granule size. Similar phenotypes were observed in salivary glands expressing *Rab11-RNAi*. To further confirm these results, we examined the localization of mucins in whole animals at the onset of metamorphosis. Mucins are expelled out of the salivary gland lumen and onto the surface of control animals at puparium formation (Biyasheva et al., 2001; Kang et al., 2017), forming a visible fluorescent ‘halo’ around the prepupa (Fig. 5D). In contrast, mucins were present only within the salivary glands and not on the surface of animals upon salivary gland-specific overexpression of *Rab11^DN^* (Fig. 5D), confirming that exocytosis fails upon loss of *Rab11* function.

**Figure 5.**
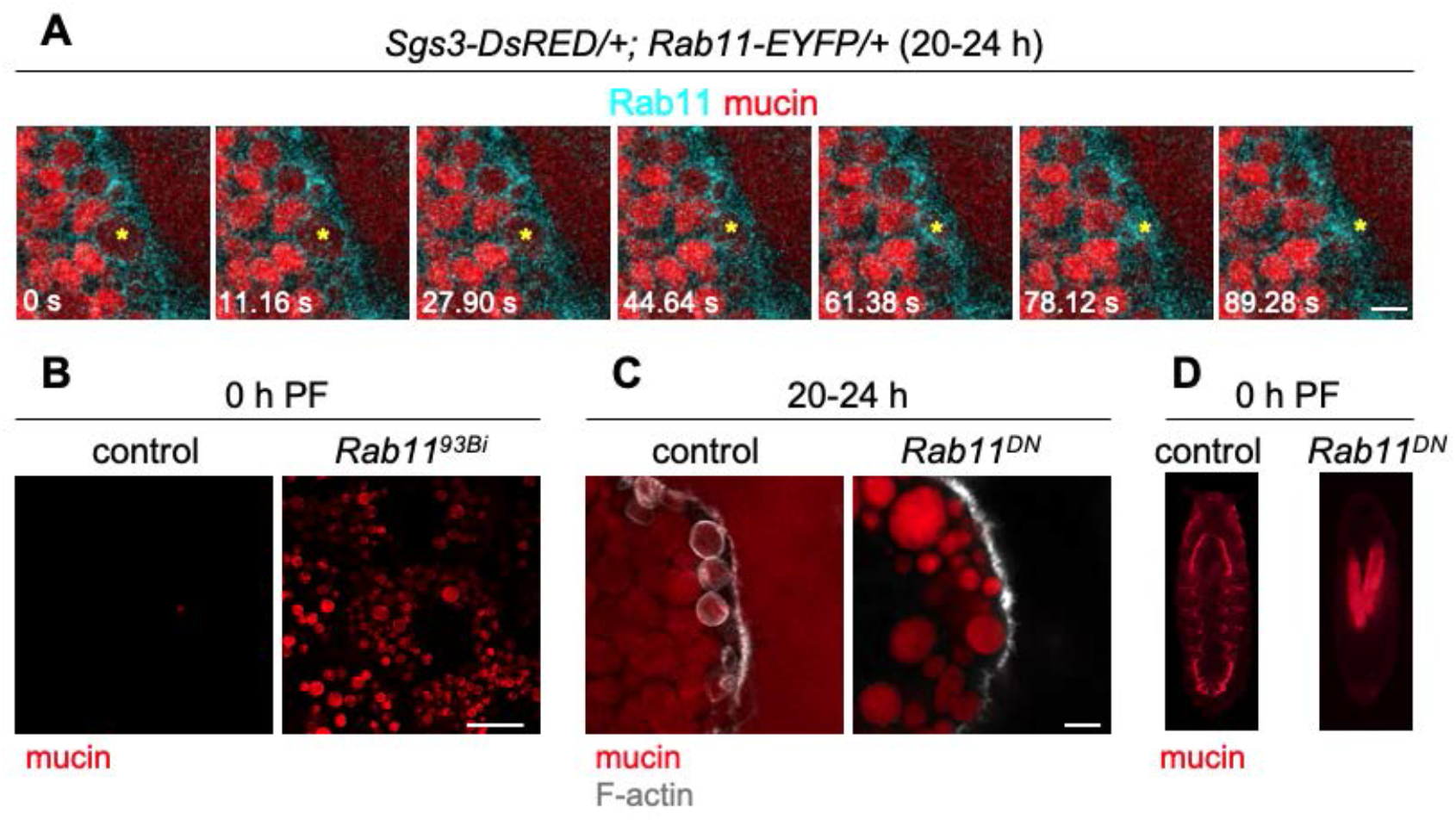
Rab11 function is required for exocytosis. **(A)** Live-cell time-lapse imaging of Rab11-EYFP (cyan) and mucin (red) localization during granule exocytosis shows that Rab11 stays associated with granule membranes during exocytosis. Yellow asterisk denotes a granule undergoing exocytosis over the course of the time-lapse. Images shown are stills from Movie 1 and represent a maximum intensity projection of nine optical slices acquired at a 0.31 μm step size; these images were not deconvolved. The salivary gland lumen is on the right side of the images. Note that this time-lapse shows only the final steps of secretory granule fusion with the plasma membrane. **(B)** Live-cell imaging of mucins (red) in control (*Sgs3-GFP/+*) and *Rab11^93Bi^* mutant (*Sgs3-GFP/+; Rab11^93Bi^/Rab11^93Bi^*) salivary glands shows that *Rab11* mutant glands have secretion defects at the onset of metamorphosis (0 h after puparium formation, PF). *Rab11^93Bi^* image shows a single slice from a z-stack comprising 31 optical slices at a 0.36 μm step size. **(C)** Phalloidin staining (gray) to detect F-actin in fixed cells shows that F-actin wraps around mucin granules (red) undergoing exocytosis in control (*Sgs3-DsRED/+; Sgs3>/+*) but not *Rab11^DN^*-expressing (*Sgs3-DsRED/+; Sgs3>/UAS-Rab11^DN^-YFP*) salivary glands at 20-24 h after Sgs3. Control image shows a single slice from a z-stack comprising eight optical slices at a 0.20 μm step size. *Rab11^DN^* image shows a single slice from a z-stack comprising 21 slices at a 0.36 μm step size. The salivary gland lumen is on the right side of the images. Note that the variability in secretory granule size observed in *Rab11^DN^*-expressing glands likely results from a delay in expression of that transgene (see text for details). **(D)** Imaging of whole puparia at the onset of metamorphosis (0 h PF) shows that mucins (red) have been expelled onto the surface of the animal in controls but remain trapped in the glands in *Rab11^DN^*-expressing glands. Genotypes are as shown in (C). Scale bars in A, C: 5 μm; B: 20 μm.

We next wanted to understand why loss of *Rab11* function inhibits secretory granule maturation and exocytosis. Previous work by us and others demonstrates that defects in secretory granule membrane fusion protein trafficking results in both maturation and exocytosis defects (Neuman and Bashirullah, 2018; Ma et al., 2020; Burgess et al., 2012), prompting us to test if these membrane fusion proteins were present on secretory granule membranes upon loss of *Rab11*. The SNARE protein SNAP-24 was present on the membranes of secretory granules in control glands (Fig. S5A), consistent with previously reported results (Niemeyer and Schwarz, 2000; Neuman and Bashirullah, 2018). SNAP-24 was also present on the membranes of small secretory granules in salivary glands with RNAi knockdown of *Rab11* (Fig. S5C), indicating that a defect in membrane fusion protein trafficking is not responsible for the maturation or exocytosis defects observed upon loss of *Rab11*.

As a next step to understanding why exocytosis fails upon loss of *Rab11*, we examined the machinery required for physical fusion of secretory granule membranes with the apical plasma membrane. Previous work has demonstrated that the actomyosin cytoskeleton is required for exocytosis of these large mucin-containing secretory granules, and assembly of filamentous actin (F-actin) is the first observable event during this process (Rousso et al., 2015; Tran et al., 2015). Consistently, we observed filamentous actin (F-actin), assessed by phalloidin staining, wrapped around granules undergoing exocytosis in control salivary glands, and the presence of mucins in the lumen indicated that other secretory granules had already undergone exocytosis (Fig. 5C). In contrast, no mucin was visible in the lumen of salivary glands overexpressing *Rab11^DN^* at the 20-24 h timepoint, and no secretory granules near the apical membrane were wrapped in F-actin (Fig. 5C). Note that the variability in secretory granule size in *Rab11^DN^*-expressing glands is likely due to the delayed onset of expression of Rab11^DN^-YFP, as described earlier. Taken together, these results suggest that Rab11 may play a functional role in promoting F-actin-dependent fusion of mucin-containing secretory granules during exocytosis.

### Small granules lacking Rab1 are secreted

Our next goal was to determine if loss of *Rab1* affected exocytosis. *Rab1* knockdown did not appear to perturb trafficking of SNAP-24, since this SNARE protein was still present on the membranes of small secretory granules (Fig. S5A, B). We also did not observe any significant secretion defects at the onset of metamorphosis in salivary glands expressing *Rab1-RNAi*. Surprisingly, unlike *Rab11^DN^*, salivary glands expressing *Rab1^DN^* had visible mucins in the lumen and tiny secretory granules near the apical membrane wrapped in F-actin at the 20-24 h timepoint (Fig. 6C). To confirm these results, we performed live-cell time-lapse imaging of mucin granules and F-actin dynamics using the fluorescently tagged F-actin binding peptide Lifeact-ruby (Riedl et al., 2008). Consistent with previous reports (Rousso et al., 2015; Tran et al., 2015), we observed F-actin assemble around mucin granules as they were undergoing exocytosis in control cells (Movie 2, Fig. 6A). Similarly, F-actin assembled around small granules in salivary glands expressing *Rab1-RNAi*, and these small granules were secreted (Movie 3, Fig. 6B). This data shows that immature granules lacking Rab1 can be secreted and suggests that an increase in mucin granule size is not required for exocytosis.

**Figure 6:**
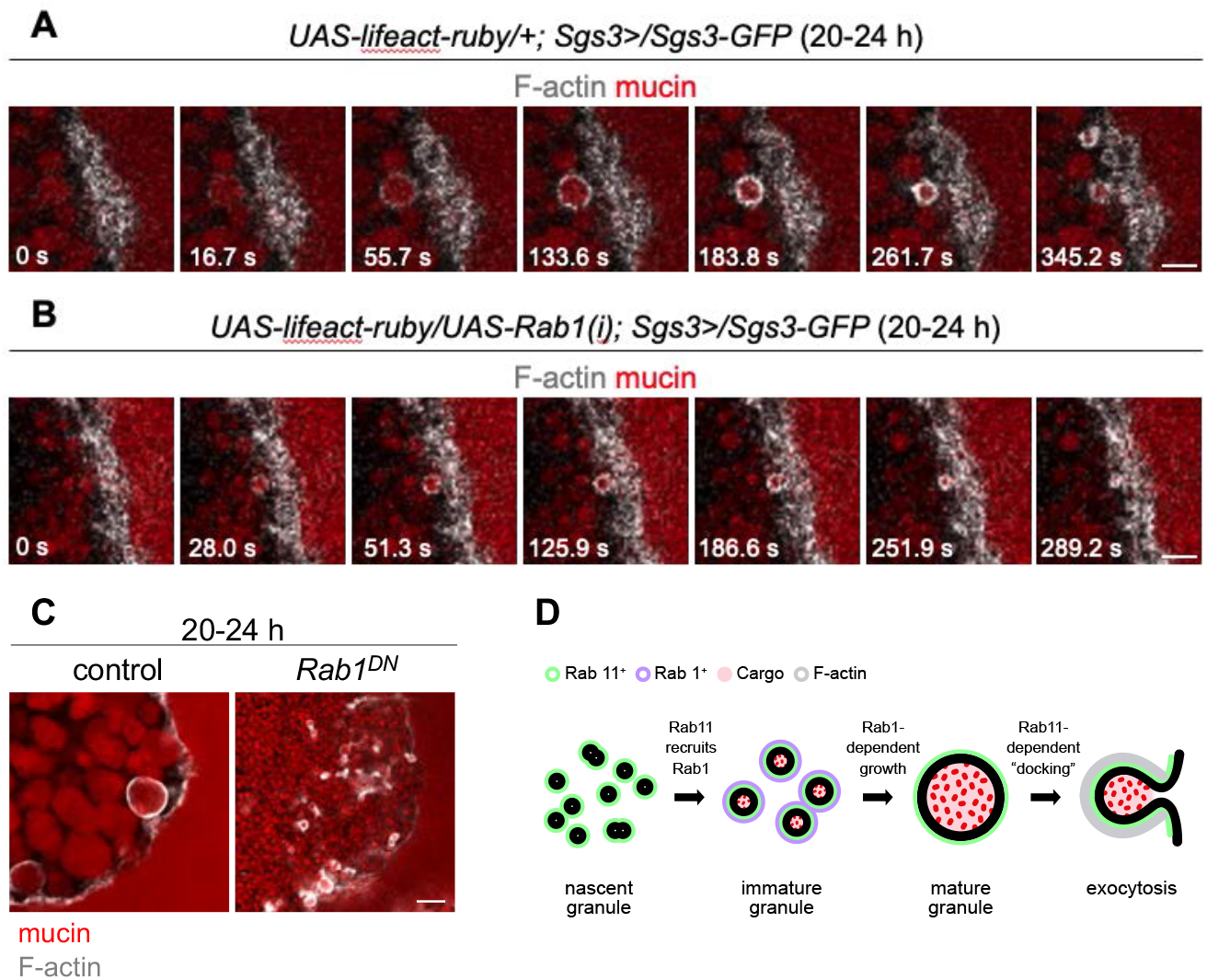
Small granules lacking Rab1 undergo exocytosis. **(A)** Live-cell time-lapse imaging of the actin-binding peptide Lifeact-ruby to detect F-actin (gray) and mucins (red) in control cells during the process of exocytosis shows that F-actin forms around granules undergoing exocytosis. Images shown are stills from Movie 2 and represent single slices from a z-stack comprising 16 slices at a 0.75 μm step size. The salivary gland lumen is on the right side of the images. **(B)** Live-cell time-lapse imaging of F-actin (gray) and mucins (red) shows that small granules in *Rab1-RNAi-* expressing cells undergo exocytosis. Images shown are stills from Movie 3 and represent single slices from a z-stack comprising 13 slices at 1.0 μm step size. The salivary gland lumen is on the right side of the images. The time-lapses in (A) and (B) show events that occur immediately prior to and during exocytosis. **(C)** Phalloidin staining to detect F-actin (gray) in fixed cells shows that F-actin wraps around normal size granules undergoing exocytosis in controls (*Sgs3-DsRED/+; Sgs3>/+*) and small granules undergoing exocytosis in *Rab1^DN^*-expressing (*Sgs3-DsRED/UAS-Rab1^DN^-YFP; Sgs3>/+*) glands at 20-24 h after Sgs3. Control image shows a single slice from a z-stack comprising 19 optical slices at a 0.36 μm step size; Rab1^DN^ image shows a single slice from a z-stack comprising 15 optical slices at a 0.20 μm step size. The salivary gland lumen is on the right side of the images. Scale bars: 5 μm. **(D)** Summary model. Mucin-containing secretory granules undergo two distinct ‘phases’ of growth; the first involves a rapid, Rab1-independent fusion of nascent granules to generate an ~18-fold increase in volume in just 4 h. The second phase of growth begins at 4 h after biogenesis and involves a more gradual Rab1-dependent ~15-fold increase in granule volume over the following 16 h. Rab11 is present on nascent granule membranes and recruits Rab1 to immature granule membranes. Rab1 is removed from mature granules, while Rab11 remains on granule membranes and regulates secretory granule “docking” to the plasma membrane and/or F-actin recruitment during exocytosis.

## DISCUSSION

Secretory granule maturation is a poorly understood process during regulated exocytosis. Here, we have identified a new role for Rab proteins in the regulation of secretory granule size during maturation. Rab11 is present on nascent secretory granule membranes and is required for recruitment of Rab1; Rab1, in turn, drives growth of immature secretory granules, and Rab1 removal signals the completion of maturation. Rab11 appears to have a second functional role in initiating secretory granule fusion with the plasma membrane during regulated exocytosis (Fig. 6D). Overall, these findings provide unexpected new insights into the function of Rab proteins during secretory granule maturation and regulated exocytosis.

Our results demonstrate that *Rab1* plays a critical functional role in promoting growth of immature granules during maturation. However, how *Rab1* facilitates fusion of immature granules remains an unanswered question. Secretory granule membrane protein trafficking appears unaffected upon loss of *Rab1*; therefore, other as-yet-unidentified factors must function downstream of *Rab1* to regulate granule-granule fusion. Rab proteins function through direct interaction with effector proteins; therefore, one or more *Rab1* effectors likely play a role in secretory granule growth. However, most of the known *Rab1* effectors localize to the ER/ERES/Golgi to regulate ER exit (Homma et al., 2021), raising the possibility that *Rab1* may work with a distinct subset of effectors that are specific to secretory granules. What regulates removal of Rab1 from mature granules also remains an open question. It is possible that removal of Rab1 may be triggered by a granule-autonomous process that ‘senses’ when the granule has reached its final mature size; alternatively, removal of Rab1 could be triggered by an independent developmental signal. Rab GAPs are frequently involved in removal of Rabs from their target membrane (Müller and Goody, 2018); therefore, a change in the localization or developmental expression levels of *Rab1* GAPs may facilitate Rab1 removal.

*Rab11* appears to have two roles during mucin exocytosis: first in recruiting Rab1 to immature granule membranes, and second in promoting mucin granule fusion with the plasma membrane during secretion. However, we cannot eliminate a possible earlier function for *Rab11* in regulating fusion of nascent granules due to technical limitations. The *Sgs3-GAL4* driver does not drive robust expression of dsRNA hairpins early enough to interfere with a function during the first growth phase of nascent granules, and the *Rab11^93Bi^* allele is quite weak, with a significant fraction of the animals surviving to adulthood (Jankovics et al., 2001). In contrast, the requirement for *Rab11* function during exocytosis, like the role of *Rab11* in recruiting Rab1 to immature granules, is much more conclusive. However, we do not yet know how *Rab11* promotes exocytosis. During mucin exocytosis, the granules fuse with the apical membrane, forming a ‘neck-like’ fusion pore; this is followed by the recruitment of F-actin and non-muscle myosin which generate a contractile force to drive cargo release (Rousso et al., 2015; Tran et al., 2015). *Rab11* could be required for either of these steps or for an earlier membrane attachment/tethering step. *Sec15*, a component of the exocyst complex that plays a role in tethering of vesicles to the plasma membrane, has been reported to be an effector of Rab11 (Wu et al., 2005; Zhang et al., 2004), suggesting that *Rab11* function may be required for plasma membrane tethering of mucin-containing vesicles in the larval salivary glands. Given the high penetrance and severity of the secretion defects observed upon loss of *Rab11* function, we posit that *Rab11* function is likely required for an early membrane attachment or fusion pore formation step during mucin exocytosis. Future studies will be required to test this model.

Secretory granule maturation, including the process of granule fusion, is thought to be essential for regulated exocytosis, as immature granules are not responsive to stimulation with secretagogues (Bonnemaison et al., 2013). However, our data indicates that small granules lacking Rab1 can still undergo regulated exocytosis, suggesting that at least some aspects of secretory granule maturation may be dispensable. Alternatively, this finding raises the possibility that Rab1 may serve as a ‘marker’ for immature granules, and the presence of Rab1 on secretory granule membranes may prevent those granules from undergoing exocytosis. Our unpublished observations suggest that the mucins secreted from these small granules are abnormal. A number of other processes are known to occur during secretory granule maturation, including refinement of secretory granule membrane and cargo content (Kögel and Gerdes, 2010a; Bonnemaison et al., 2013), suggesting that *Rab1* function may be required for other processes during secretory granule maturation beyond granule growth and that the presence of Rab1 on granules may inhibit the precocious release of immature granules.

One of the unique features of mucin granules in the larval salivary glands is their large size; mature granules reach an average volume of 30 μm^3^. This large size presents significant mechanical challenges for membrane fusion events, suggesting that these granules may require additional factors to generate sufficient contractile forces to drive granule-granule fusion during maturation and granule-plasma membrane fusion during exocytosis. However, large secretory granules are not unique to the *Drosophila* larval salivary glands; several mammalian cell types also produce and secrete large vesicles, including digestive enzymes in the exocrine pancreas (Geron et al., 2013), surfactants in the lungs (Miklavc et al., 2015), and von Willebrand factor in endothelial cells (Nightingale et al., 2011). Future analysis of the localization and function of Rab1 and Rab11 in these systems, as well as in other professional secretory cells, may reveal a shared requirement for these proteins in secretory vesicle trafficking.

## MATERIALS AND METHODS

### Fly stocks and husbandry

The following fly stocks were obtained from the Bloomington *Drosophila* Stock Center: all *UAS-Rab-RNAi* lines (stock identifiers listed in Table S1), *É^1118^, Sgs3-GFP, Rab1-EFYP, Rab3-EFYP, Rab5-EYFP, Rab10-EFYP, Rab11-EYFP, Rab27-EFYP, UAS-Rab1^DN^-YFP, UAS-Rab11^DN^-YFP, Sgs3-GAL4, UAS-lifeact-ruby, Rab11^93Bi^, UAS-Sec31-RFP, UAS-Golgi-RFP. Sgs3-DsRED* was provided by A. Andres (University of Nevada, Las Vegas, NV, USA). We have used the “>“ symbol in genotypes as shorthand for “GAL4.” All experimental crosses were grown in uncrowded vials or bottles on standard cornmeal-molasses media in an incubator set to 25°C.

### Developmental staging

Animals were synchronized at the onset of mucin/glue biogenesis as previously described (Neuman et al., 2021). Given that there is some degree of variability in developmental rates between animals, a second layer of tissue-autonomous synchronization was used to further confirm developmental stages. Since glue protein synthesis begins in the cells at the distal tip of the glands and progresses forward (Biyasheva et al., 2001), the number of cells that synthesized glue was used to confirm developmental timing according to the following criteria: 0-4 h after Sgs3- glue protein visible only in the cells at the distal tip of the glands; 4-8 h after Sgs3- glue protein visible in the distal one-third to one-half of the gland; 8-12 h after Sgs3- glue protein visible in the distal one-half to two-thirds of the gland; 12-16 h after Sgs3- glue protein visible in all the large acinar cells of the gland; 16-20 h after Sgs3- glue protein visible in all cells of the gland, including smaller acinar cells located near the salivary gland duct; 20-24 h after Sgs3- onset of secretion with glue protein visible in the lumen. Only cells near the distal tip of the gland were imaged to ensure consistency across developmental stages and genotypes.

### Quantitative real-time PCR (qPCR)

qPCR was performed as previously described (Ihry et al., 2012). Total RNA was isolated from dissected salivary glands of the appropriate developmental stage and genotype using the RNeasy Plus Mini Kit (Qiagen). 400 ng of total RNA was used to synthesize cDNA with the SuperScript III First-Strand Synthesis System (Invitrogen). Samples were collected and analyzed in biological triplicate. qPCR was performed on a Roche LightCycler 480 with LightCycler 480 SYBR Green I Master Mix (Roche). Relative Expression Software Tool (REST) was used to calculate relative expression and *p*-values (Pfaffl et al., 2002). Expression was normalized to the reference gene *rp49;* primers for *rp49* were previously published (Ihry et al., 2012). New primers for *Rab1* and *Rab11* were designed using FlyPrimerBank (Hu et al., 2013). *Rab1* F: 5′-CGA CGG AAA GAC CAT TAA ACT GC-3′; *Rab1* R: 5′- GCG CCC CTA TAA TAT GAA GAC G-3′; *Rab11* F: 5′- ATT TGC TCT CAC GTT TCA CGC-3′; *Rab11* R: 5′- GCC ATC GAC CTC TAT GCT GC-3′.

### Confocal microscopy and immunofluorescence

All images except those in Fig. 5C, 6C, and S5 were obtained from live, unfixed tissue. For live tissue imaging, salivary glands of the appropriate developmental stage and genotype were dissected in PBS and mounted in 1% low-melt agarose (Apex Chemicals) made in PBS. Tissues were imaged for no more than 15 min after mounting, and imaging was carried out at room temperature. At least 10 salivary glands were imaged per experiment. Images were acquired using an Olympus FV3000 laser scanning confocal microscope (60x oil immersion objective, NA 1.42 or 100x oil immersion objective, NA 1.49) with FV31S-SW software. Brightness and contrast were adjusted post-acquisition with FV31S-SW software. Images acquired as z-stacks (as indicated in figure legends) were deconvolved using three iterations of the Olympus CellSens Deconvolution for Laser Scanning Confocal Advanced Maximum Likelihood algorithm. Movies were acquired using the Olympus FV3000 resonant scan head. For fixed tissue imaging, immunofluorescent staining for SNAP-24 was carried out as previously described (Yin et al., 2007; Neuman and Bashirullah, 2018). Salivary glands of the appropriate stage and genotype were dissected in PBS, fixed for 30 min at room temperature in PBS with 0.1% Triton-X 100 (PBST)/4% formaldehyde and blocked overnight at 4°C in PBST/4% BSA. The SNAP-24 antibody (gift from T. Schwarz, Harvard Medical School) (Niemeyer and Schwarz, 2000) was diluted at 1:200 in PBST/4% BSA and tissues were stained overnight at 4°C. α-rabbit Cy3 (Jackson Immuno-Research Labs 711-165-152) secondary antibody was diluted at 1:200 in PBST/4% BSA. For phalloidin staining, salivary glands of the appropriate stage and genotype were fixed and blocked as described above. Tissues were incubated in 1:250 Oregon Green 488 Phalloidin (Invitrogen O7466) for 30 min at room temperature. Stained tissues were mounted in Vectashield (Vector Laboratories) and imaged as described above. Quantification of granule volume was done in ImageJ, and z-stack data was used to confirm that the diameter was measured at the medial cross-section of each granule. Box and whisker plot was generated using Prism. Quantification of the percentage of granules with Rab1-EFYP was done using FV31S-SW software. Images of whole puparia were obtained using an Olympus SXZ16 stereomicroscope coupled to an Olympus DP72 digital camera with DP2-BSW software.

## ACKNOWLEDGEMENTS

Stocks obtained from the Bloomington *Drosophila* Stock Center (NIH P40OD018537) were used in this study. The authors thank Lora Luo for assistance with illustration.

## COMPETING INTERESTS

The authors declare no competing interests.

## FUNDING

This work was supported in part by the National Institutes of Health (GM123204 to A.B.).

## AUTHOR CONTRIBUTIONS

Conceptualization: S.D.N., A.B; Methodology: S.D.N., A.B.; Validation: S.D.N., A.B.; Investigation: S.D.N., A.R.L., J.E.S., A.T.C.; Data curation: S.D.N., A.R.L.; Writing-original draft: S.D.N.; Writing-review and editing: S.D.N., A.B.; Visualization: S.D.N., A.B.; Supervision: A.B.; Project administration: A.B.; Funding acquisition: A.B.

---

**Table S1.** Summary of *Rab-RNAi* screen results. All RNAi lines were crossed to flies containing the *Sgs3-GFP* and *Sgs3-GAL4* transgenes. ++ indicates a strong reduction in granule size; + indicates a moderate reduction in granule size; - indicates no size defect. Highlighted stocks were validated for the efficiency of RNAi knockdown via qPCR and used for all detailed experimental analysis (see Fig. S1).

| Rab protein | Stock Identifier | Small granules? |
| --- | --- | --- |
| Rab1 | RRID:BDSC_27299 | ++ |
|  | RRID:BDSC_34670 | ++ |
| Rab2 | RRID:BDSC_28701 | - |
|  | RRID:BDSC_34922 | - |
| Rab3 | RRID:BDSC_31691 | + |
|  | RRID:BDSC_34655 | - |
| Rab4 | RRID:BDSC_33757 | - |
| Rab5 | RRID:BDSC_30518 | + |
|  | RRID:BDSC_34832 | - |
|  | RRID:BDSC_51847 | - |
|  | RRID:BDSC_67877 | - |
| Rab6 | RRID:BDSC_27490 | - |
|  | RRID:BDSC_35744 | - |
| Rab7 | RRID:BDSC_27051 | - |
| Rab8 | RRID:BDSC_27519 | - |
|  | RRID:BDSC_34373 | - |
|  | RRID:BDSC_76059 | - |
| Rab9 | RRID:BDSC_31688 | - |
|  | RRID:BDSC_42942 | - |
| Rab9Db | RRID:BDSC_38269 | - |
| Rab9Fb | RRID:BDSC_34374 | - |
| Rab10 | RRID:BDSC_26289 | + |
| Rab11 | RRID:BDSC_27730 | + |
|  | RRID:BDSC_42709 | ++ |
| Rab14 | RRID:BDSC_28708 | - |
|  | RRID:BDSC_34654 | - |
| Rab18 | RRID:BDSC_27665 | - |
|  | RRID:BDSC_34734 | - |
| Rab19 | RRID:BDSC_34607 | - |
| Rab21 | RRID:BDSC_29403 | - |
| Rab23 | RRID:BDSC_28025 | - |
|  | RRID:BDSC_36091 | - |
|  | RRID:BDSC_55352 | - |
|  | RRID:BDSC_63689 | - |
| Rab26 | RRID:BDSC_31177 | - |
| Rab27 | RRID:BDSC_31887 | + |
|  | RRID:BDSC_35774 | - |
|  | RRID:BDSC_50537 | - |
| Rab30 | RRID:BDSC_31120 | - |
| Rab32 | RRID:BDSC_28002 | - |
|  | RRID:BDSC_38956 | - |
| Rab35 | RRID:BDSC_28342 | - |
|  | RRID:BDSC_67952 | - |
|  | RRID:BDSC_80457 | - |
| Rab39 | RRID:BDSC_25953 | - |
|  | RRID:BDSC_51689 | - |
| Rab40 | RRID:BDSC_29579 | - |
|  | RRID:BDSC_61308 | - |
|  | RRID:BDSC_80472 | - |
| RabX1 | RRID:BDSC_28033 | - |
| RabX2 | RRID:BDSC_32360 | - |
|  | RRID:BDSC_53928 | - |
| RabX4 | RRID:BDSC_28704 | - |
|  | RRID:BDSC_44070 | - |
| RabX5 | RRID:BDSC_28045 | - |
| RabX6 | RRID:BDSC_26281 | - |
|  | RRID:BDSC_53252 | - |

**Figure S1.**
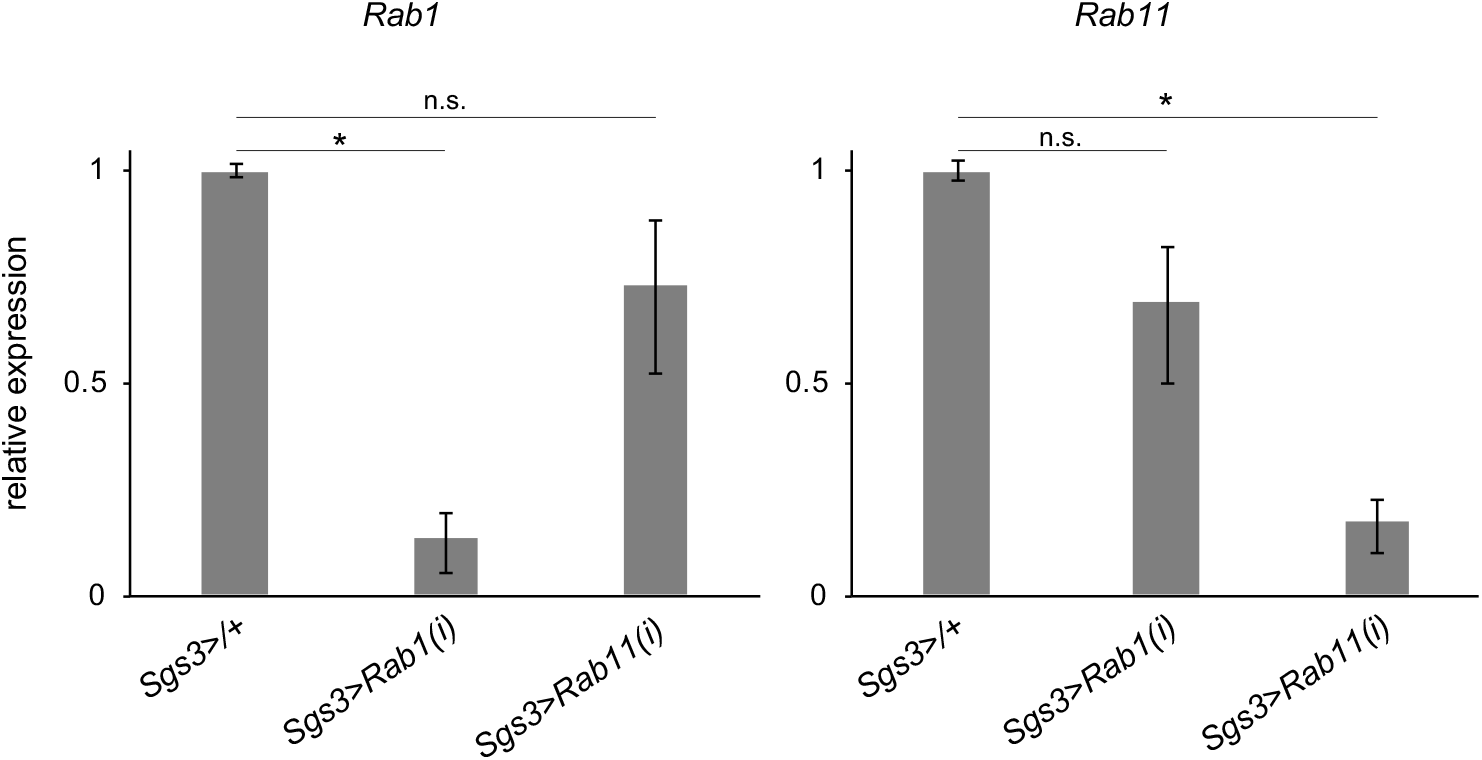
Validation of RNAi knockdown efficiency. qPCR analysis of *Rab1* and *Rab11* expression levels in dissected salivary glands from animals synchronized at the onset of metamorphosis (0 h after puparium formation, PF) shows that *Rab1-RNAi* knocks down *Rab1* expression levels but does not significantly affect *Rab11* expression levels. Similarly, *Rab11-RNAi* reduces *Rab11* expression levels but does not significantly alter *Rab1* expression levels. *y*-axis shows relative expression and *x*-axis shows the genotypes analyzed. Expression was normalized to *rp49* and samples were analyzed in biological triplicate. Error bars and statistics were determined using REST analysis. Asterisks indicate *p*>0.05.

**Figure S2.**
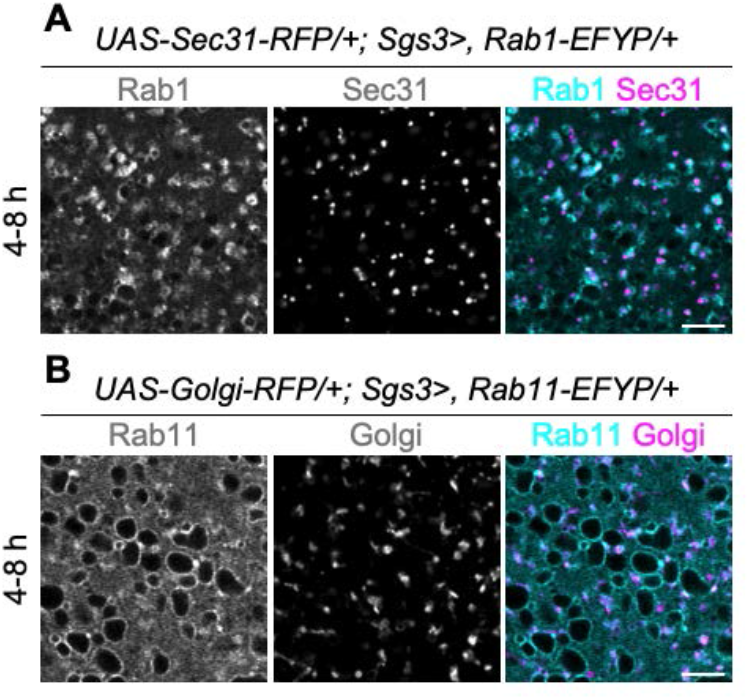
Rab1 also localizes to ER exit sites and Rab11 also localizes to Golgi. **(A)** Live-cell imaging of Rab1-EFYP (cyan) and Sec31-RFP (magenta), a component of the COPII coat complex and marker for ER exit sites (ERES), shows that Rab1 localizes to ERES in addition to localizing to secretory granule membranes at 4-8 h after Sgs3. Image shown is a single slice from a z-stack comprising ten optical slices at a 0.31 μm step size. **(B)** Live-cell imaging of Rab11-EFYP (cyan) and the pan-Golgi marker Golgi-RFP (magenta), shows that Rab1 localizes to the Golgi in addition to localizing to secretory granule membranes at 4-8 h after Sgs3. Note that small Rab11-positive puncta that do not co-localize with Golgi-RFP are also present; these are likely recycling endosomes. Image shown is a single slice from a z-stack comprising 17 optical slices at a 0.30 μm step size. Scale bars: 5 μm.

**Figure S3.**
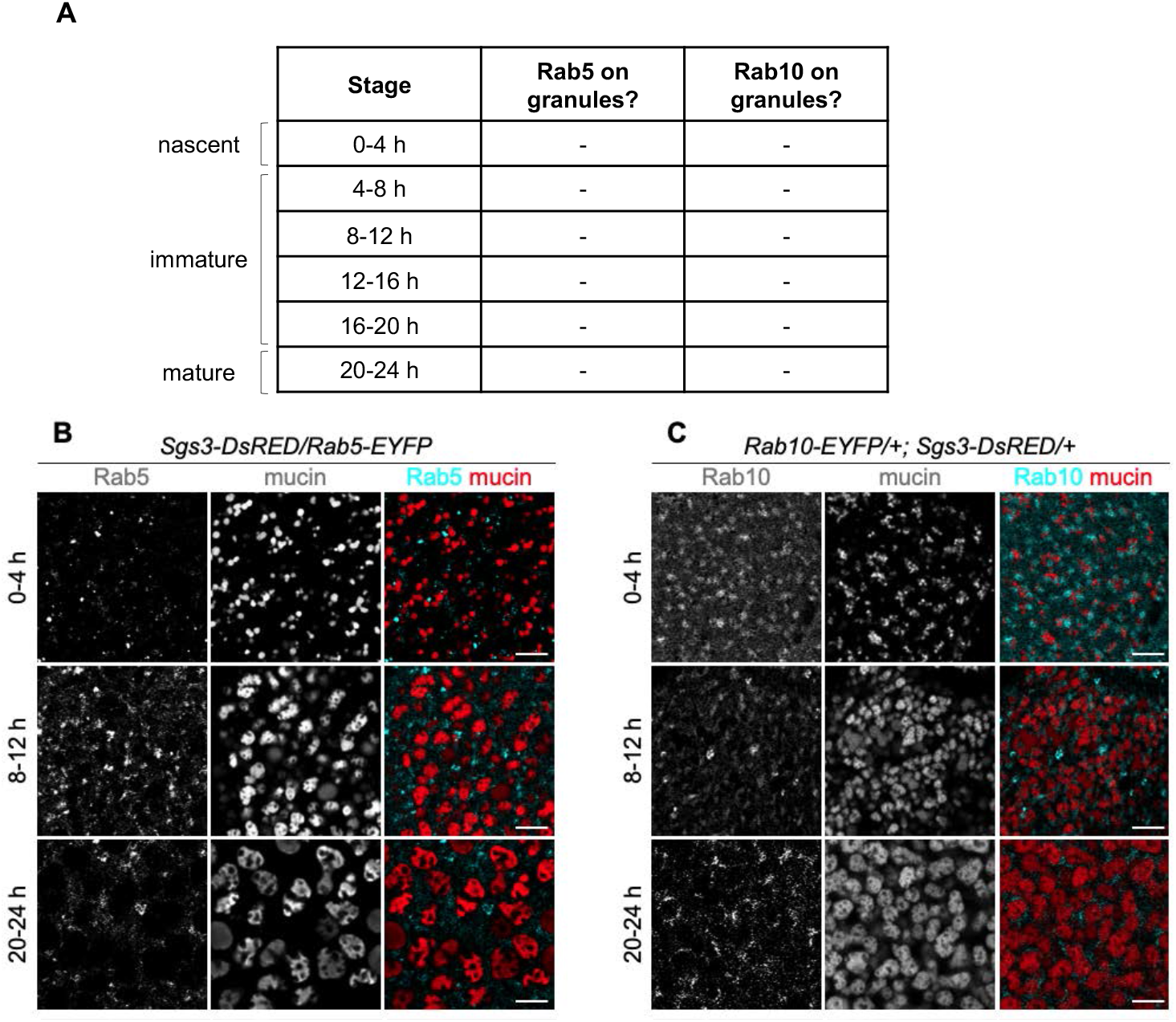
Rab5 and Rab10 do not localize to mucin granule membranes. **(A)** Summary table showing the presence of Rab5 and Rab10 on secretory granule membranes in staged salivary glands. Neither protein is present on secretory granule membranes at any of the developmental stages analyzed. Developmental stages are shown in h after Sgs3. *n*=10 glands imaged per timepoint. **(B)** Live-cell imaging analysis of Rab5-EYFP (cyan) localization with mucins (red). Rab5 is absent from granule membranes but localizes in small puncta that are likely early endosomes (Zhang et al., 2007; Dunst et al., 2015) at 0-4, 8-12, and 20-24 h after Sgs3. Images shown are a single slice from a z-stack comprising three slices (0-4 h), four slices (8-12 h), or five slices (20-24 h) at a 0.31 μm step size. **(C)** Live-cell imaging analysis of Rab10-EYFP (cyan) localization with mucins (red). Rab10 is absent from granule membranes but localizes in small puncta that are likely recycling endosomes (Chan et al., 2011; Gaudet et al., 2011) at 0-4, 8-12, and 20-24 h after Sgs3. Images shown are a single slice from a z-stack comprising 16 slices (0-4 h), 13 slices (8-12 h) or six slices (20-24 h) at a 0.31 μm step size. Scale bars: 5 μm. Note that we did not observe substantial expression of Rab3- or Rab27-EFYP in salivary glands; therefore, these proteins were not pursued further.

**Figure S4.**
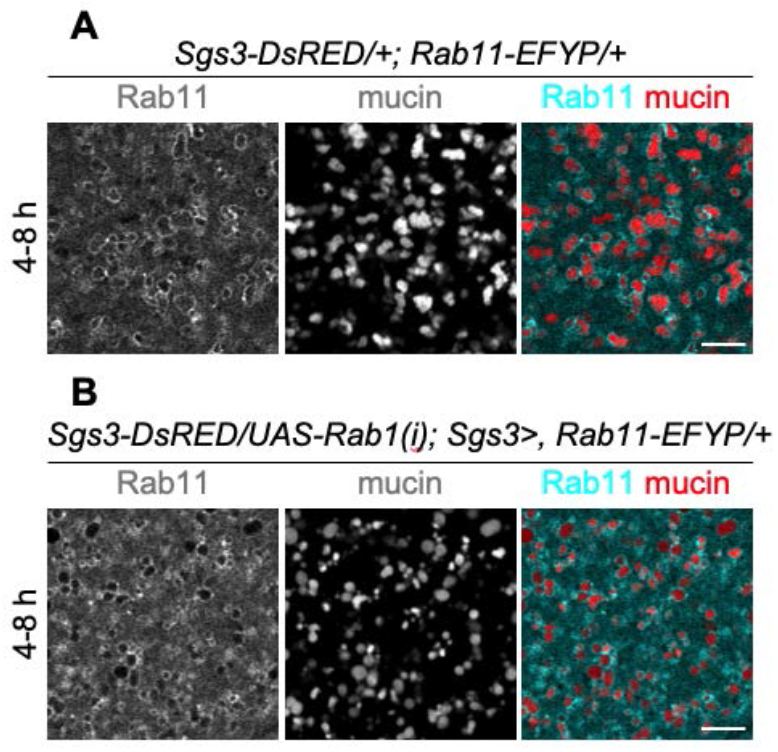
*Rab1* knockdown does not affect Rab11 localization. **(A)** Live-cell imaging of Rab11-EFYP (cyan) and mucins (red) shows that Rab11 is present on mucin granule membranes at 4-8 h after Sgs3. Image shown is a single slice from a z-stack comprising ten optical slices at a 0.31 μm step size. **(B)** Live-cell imaging of Rab11-EYFP (cyan) localization with mucins (red) in salivary glands expressing *Rab1-RNAi* shows that Rab11 is still present around secretory granule membranes upon knockdown of *Rab1* expression at 4-8 h after Sgs3. Image shown is a single slice from a z-stack comprising ten optical slices at a 0.31 μm step size. Scale bars: 5 μm.

**Figure S5.**
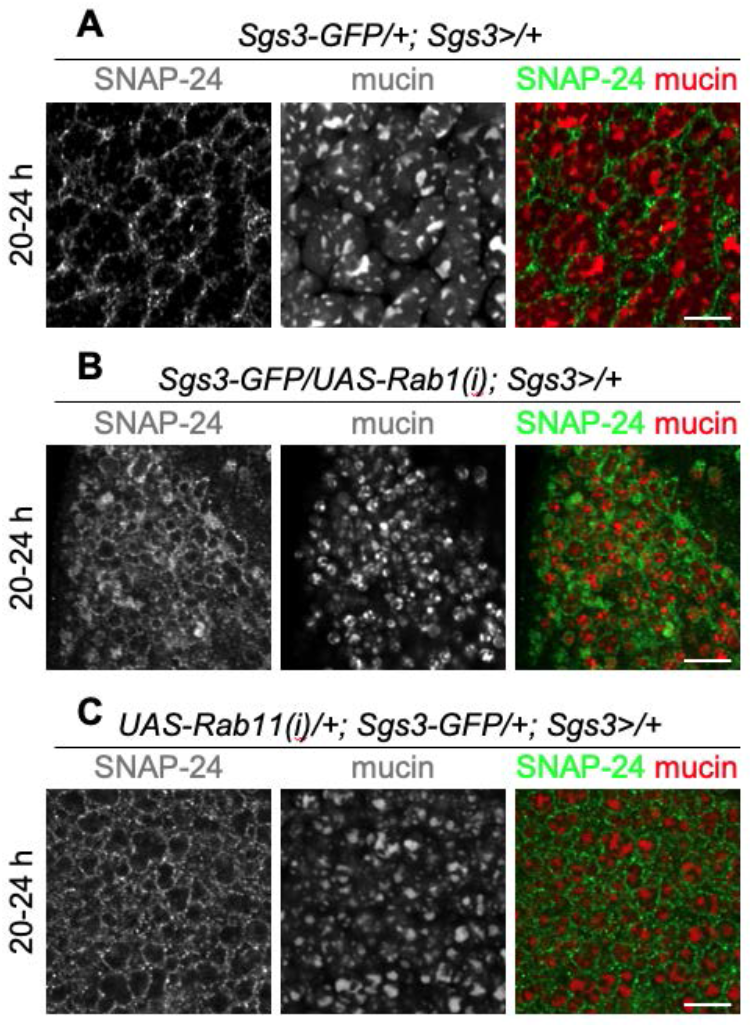
Knockdown of *Rab1* or *Rab11* does not affect secretory granule membrane protein localization. **(A)** Antibody staining for the SNARE protein SNAP-24 (green) in fixed cells shows that this secretory granule membrane protein is present around mucin granules (red) in control salivary glands at 20-24 h after Sgs3. Image shown is a single slice from a z-stack comprising 13 optical slices at a 0.37 μm step size. **(B)** SNAP-24 (green) is present around small mucin granules (red) in *Rab1-RNAi*-expressing cells at 20-24 h after Sgs3. Image shown is a single slice from a z-stack comprising nine optical slices at 0.37 μm step size. **(C)** SNAP-24 (green) is present around small mucin granules (red) in *Rab11-RNAi*-expressing cells at 20-24 h after Sgs3. Image shown is a single slice from a z-stack comprising 13 optical slices at 0.37 μm step size. Scale bars: 5 μm.

**Movie 1. Rab11 is present on secretory granule membranes during exocytosis.** Live-cell time-lapse imaging of Rab11-EYFP (cyan) and mucins (red) shows that Rab11 stays associated with secretory granule membranes and is deposited on the apical plasma membrane during exocytosis. Arrow points to a granule undergoing exocytosis. Movie shows a maximum intensity projection of nine optical slices acquired at a 0.31 um step size and was not deconvolved. A total of 100 frames were captured, comprising ~9 mins total time. Full genotype: *Sgs3-DsRED/+; Rab11-EYFP/+*.

**Movie 2. F-actin dynamics during exocytosis in control cells.** Live-cell time-lapse imaging of Lifeact-ruby (gray) to visualize F-actin and mucins (red) during exocytosis in control cells shows that F-actin forms around secreting granules and drives exocytosis. Arrow points to a granule undergoing exocytosis. Movie shows a single slice from a z-stack comprising 16 total optical slices at a 0.75 μm step size. A total of 150 frames were captured, comprising ~14 mins total time. Movie shows 79 frames comprising a total of ~7.3 mins total time. Full genotype: *UAS-lifeact-ruby/+; Sgs3>/Sgs3-GFP*.

**Movie 3. Small granules lacking Rab1 can be secreted.** Live-cell time-lapse imaging of Lifeact-ruby (gray) to visualize Factin and mucins (red) during exocytosis in *Rab1-RNAi-expressing* cells shows that small granules undergo exocytosis. Arrow points to a granule undergoing exocytosis. Movie shows a single slice from a z-stack comprising 13 optical slices at a 1.0 μm step size. A total of 200 frames were captured, comprising ~15 mins total time. Movie shows 69 frames comprising a total of ~5.4 mins total time. Full genotype: *UAS-lifeact-ruby/UAS-Rab1-RNAi; Sgs3>/Sgs3-GFP*.

